# Myxoma virus lacking the host range determinant M062 stimulates cGAS-dependent type 1 interferon response and unique transcriptomic changes in human macrophages

**DOI:** 10.1101/2022.01.30.478420

**Authors:** Steven J Conrad, Erich A Peterson, Jason Liem, Richard Connor, Bernice Nounamo, Martin Cannon, Jia Liu

## Abstract

The evolutionarily successful poxviruses possess effective and diverse strategies to circumvent or overcome host defense mechanisms. Poxviruses encode many immunoregulatory proteins to evade host immunity to establish a productive infection and have unique means of inhibiting DNA sensing-dependent type 1 interferon (IFN-I) responses, a necessity given their dsDNA genome and exclusively cytoplasmic life cycle. We found that the key DNA sensing inhibition by poxvirus infection was dominant during the early stage of poxvirus infection before DNA replication. In an effort to identify the poxvirus gene products which subdue the antiviral proinflammatory responses (e.g., IFN-I response), we investigated the function of one early gene that is the known host range determinant from the highly conserved poxvirus host range *C7L* superfamily, myxoma virus (MYXV) M062. Host range factors are unique features of poxviruses that determine the species and cell type tropism. Almost all sequenced mammalian poxviruses retain at least one homologue of the poxvirus host range *C7L* superfamily. In MYXV, a rabbit-specific poxvirus, the dominant and broad-spectrum host range determinant of the *C7L* superfamily is the *M062R* gene. The *M062R* gene product is essential for MYXV infection in almost all cells tested from different mammalian species and specifically inhibits the function of host <u>S</u>terile <u>α M</u>otif <u>D</u>omain-containing 9 (SAMD9), as *M062R*-null (Δ*M062R*) MYXV causes abortive infection in a SAMD9-dependent manner. In this study we investigated the immunostimulatory property of the Δ*M062R*. We found that the replication-defective Δ*M062R* activated host DNA sensing pathway during infection in a cGAS-dependent fashion and that knocking down SAMD9 expression attenuated proinflammatory responses. Moreover, transcriptomic analyses showed a unique feature of the host gene expression landscape that is different from the dsDNA-stimulated inflammatory state. This study establishes a link between the anti-neoplastic function of SAMD9 and the regulation of innate immune responses.

**Author Summary:** Poxviruses encode a group of genes called host range determinants to maintain or expand their host tropism. The mechanism by which many viral host range factors function remains elusive. Some host range factors possess immunoregulatory functions responsible for evading or subduing host immune defense mechanisms. Most known immunoregulatory proteins encoded by poxviruses are dispensable for viral replication *in vitro*. The uniqueness of MYXV *M062R* is that it is essential for viral infection in vitro and belongs to one of the most conserved poxvirus host range families, the *C7L* superfamily. There is one known host target of the MYXV M062 protein, SAMD9. SAMD9 is constitutively expressed in mammalian cells and exclusively present in the cytoplasm and has an anti-neoplastic function. Humans with deleterious mutations in SAMD9 present disease that ranges from lethality at a young age to a predisposition to myelodysplastic syndromes (MDS) that often require bone marrow transplantation. More importantly, SAMD9 serves as an important antiviral intrinsic molecule to many viruses. The cellular function of SAMD9 remains unclear mostly due to the difficulty of studying this protein, i.e., its large size, long half-life, and its constitutive expression in most cells. In this study we used M062R-null MYXV as a tool to study SAMD9 function and report a functional link between SAMD9 and the regulation of the proinflammatory responses triggered by cGAS-dependent DNA sensing.

## Introduction

Mammalian hosts have sophisticated regulation for the triggering of pro-inflammatory responses, especially after detecting danger signals in the cytoplasm. Many fundamental sensing instruments and their direct downstream signaling axes have been described, such as cGAS/STING/IRF3 axis for DNA sensing (1-3) and RNA sensing pathways (4) e.g., the RIG-I/MAVS/IRF3 axis (5). Additional factors may fine-tune the consequences of dangerous stimulus (e.g., DNA substrates) resulting in distinctive overall cellular and immunological responses. The outcome may also be tissue- and cell-type dependent. We are particularly interested in understanding the immunoregulatory mechanism of monocytes/macrophages. These immune cells are among the first responders to viral infection and also important in the maintenance of immunological microenvironment, such as the tumor environment.

Cytoplasmic surveillance for the presence of DNA is an important task for mammalian cells, as the appearance of DNA in this privileged compartment signals grave danger for the well-being of the cell. Poxviruses inhibit DNA sensing at an early time during infection and poxviruses from different genera have evolved unique strategies to circumvent host surveillance against cytoplasmic DNA. Here we investigated the phenomenon associated with transcriptomic landscape remodeling in macrophages triggered by a mutant MYXV, Δ*M062R*, and found it distinct from the classic DNA sensing which triggers the type 1 interferon (IFN-I) response. Interestingly, Δ*M062R* induced proinflammatory effect is also regulated by another host protein, SAMD9. This unusual inflammatory response induced by Δ*M062R* may explain the immunotherapeutic benefit we have observed when using Δ*M062R* in the tumor environment (6) and when treating tumor associated macrophages (TAMs) from human patients (unpublished data).

Poxviruses are exemplary probing tools in our quest to understand the host immune response(s) at the molecular and pathogenesis levels. Poxvirus must be able to evade host surveillance against cytoplasmic DNA due to their exclusive cytoplasmic life cycle to replicate their dsDNA genome. It is not surprising that many genes encoded by poxviruses antagonize DNA sensing-stimulated antiviral immune responses. In this study we utilized MYXV, a rabbit-specific poxvirus, to investigate novel host regulation of innate immune responses using a viral protein, M062, as a probing tool. Viral M062 protein is essential for MYXV infection with a classic role as host range determinant. M062 has a known host target, SAMD9, and inhibition of SAMD9 is required for a productive viral infection (7, 8). We observed that infection by the replication-defective Δ*M062R* in monocytes/macrophages led to IFN-I induction and the production of pro-inflammatory cytokines/chemokines. This proinflammatory response associated host gene expression is IRF-dependent and is regulated through DNA sensing by cGAS. The pro-inflammatory responses caused by Δ*M062R* are also regulated by SAMD9, the direct target of viral M062 protein. Interestingly, this dual regulation leads to a unique transcriptomic landscape distinct from that is induced by DNA sensing alone. We thus concluded that this additional regulation of the cGAS DNA sensing pathway through SAMD9 may act to fine-tune the consequences of DNA sensing. This finding elucidates the immunoregulatory function of SAMD9 in addition to its anti-neoplastic property and may explain the role of SAMD9 in host defense against dangerous signals.

## Results

### Early poxvirus proteins play dominant roles in suppressing dsDNA-stimulated IFN-I

Poxviruses, especially virulent poxviruses, must inhibit the host DNA sensing pathway and IFN-I production to be able to effectively replicate and spread to other cells (9). Many orthopoxviruses encode a poxvirus immune nuclease (poxin), an early poxvirus gene, for cGAS-STING-specific immune evasion (10-13), but poxviruses from non-orthopoxvirus genera may not possess such genes in their genomes, and one example is MYXV. It was reported previously that the gene product of *F17R*, a late gene, from vaccinia virus (VACV) was important for inhibiting DNA sensing (14, 15) and a homolog of the VACV *F17R* is present in MYXV, *M026R* (16). Our early experiment aimed to rule out the possibility that any early viral genes from MYXV might play a role in the inhibition of DNA sensing with VACV WR as a control. Using a well-established IRF-dependent luciferase system in macrophages in the presence of a DNA replication inhibitor, cytosine arabinoside (AraC), we found MYXV infection at the early state could already potently inhibit dsDNA-stimulated luciferase expression comparably to the ability by VACV (**Figure 1A**). The levels of inhibition in the presence of AraC from both VACV and MYXV are similar to those caused by corresponding viruses without AraC treatment (**Figure 1A**). Thus, additional viral factors that are expressed during early gene expression play dominant roles in the suppression of DNA-induced IFN-I induction in MYXV. Next, we found that infection by a replication defective virus with the essential host range gene *M062R* deleted, Δ*M062R*, lost the ability to inhibit dsDNA-stimulated IFN-I induction (**Figure 1B**). It is known that Δ*M062R* infection retains other early viral gene expression comparable to what is seen in the wildtype MYXV infection (7).

**Figure 1.**
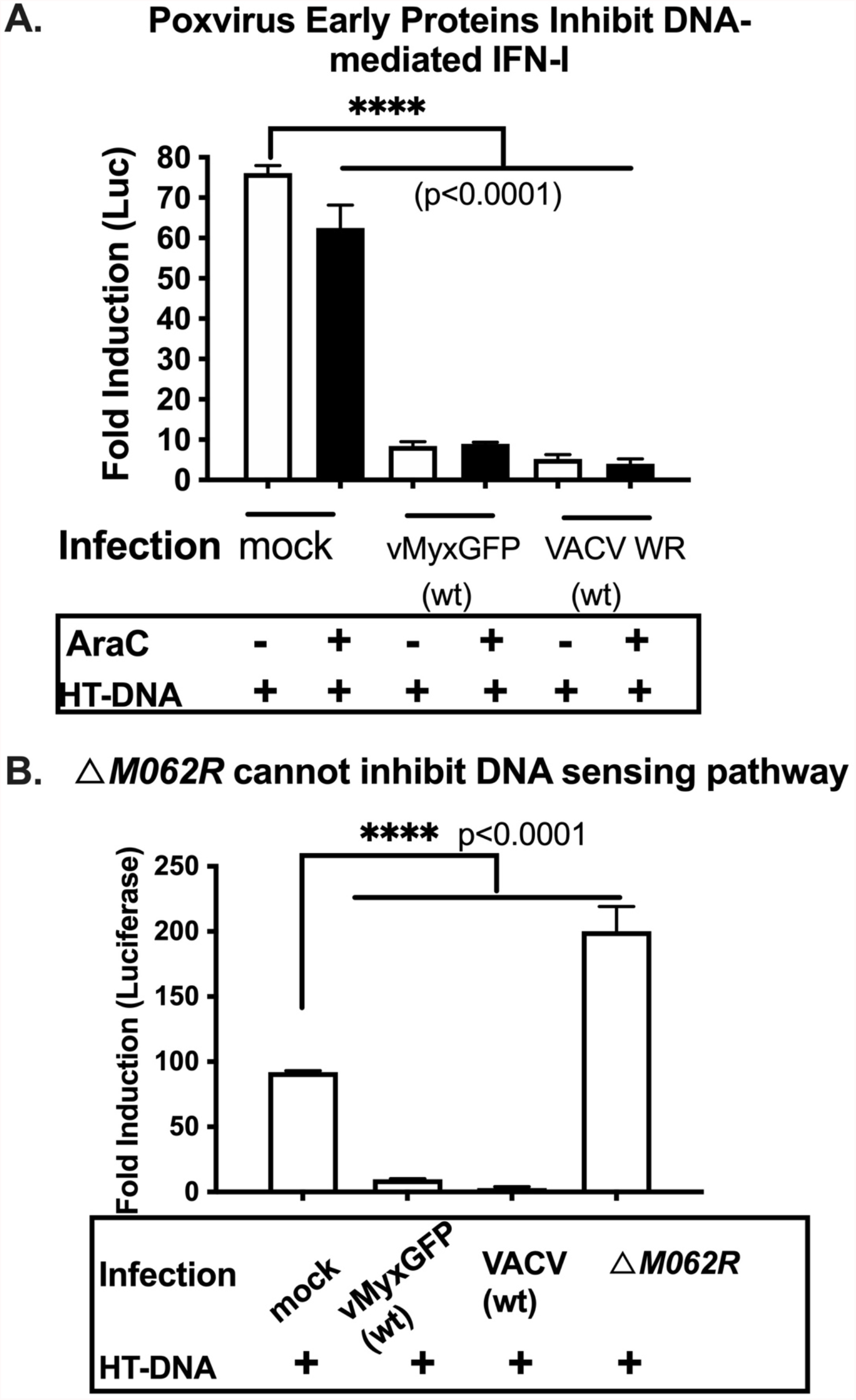
Myxoma virus *M062R* gene is important for viral inhibition of DNA sensing. **A**. Early gene expression from either wildtype MYXV or VACV inhibits dsDNA-stimulated IFN-I induction. The IRF-dependent luciferase-expressing THP-1 cell line is a surrogate system for IFN-I induction. After cells were differentiated into macrophages, viral infection was performed at an moi of 10 in the presence of AraC (100 µM) before cells were transfected with HT-DNA. In the absence of post-replicative gene products due to AraC treatment, wildtype MYXV or VACV infection significantly inhibited dsDNA-stimulated luciferase expression. Statistical analyses were performed with ordinary one-way ANOVA followed by Tukey’s multiple comparison test and p<0.05 is defined as being statically significant (****p<0.0001). Shown is a representative result of 2 biological replicates and each data point is an average of triplicate measurements (technical replicates). **B**. In the absence of *M062R* gene, the resulting ΔM*062R* MYXV loses the ability to inhibit HT-DNA stimulated luciferase expression. The same luciferase-expressing THP-1 cells were differentiated into macrophages as in “**A**”. After viral infection at a high moi of 10 with VACV or MYXV, cells were transfected with HT-DNA for 18 hrs. Supernatant was then collected to measure the luciferase activities. *M062R*-knockout MYXV could no longer inhibit dsDNA-stimulated IRF-dependent luciferase expression. Statistical analyses were performed using ordinary one-way ANOVA followed by Dunnett multiple comparison test and p<0.05 is defined as being statistically significant (****p<0.0001). Shown is a representative of 3 biological replicates and each data point is the average of 2 replicating measurements.

### *M062R*-null MYXV (Δ*M062R*) infection stimulates proinflammatory cytokine production

The MYXV early gene, *M062R*, is a broad-spectrum host range determinant from the poxvirus host range *C7L* superfamily (17) and is essential for MYXV infection (7, 8). The Δ*M062R* MYXV has a tropism defect and causes an abortive infection in almost all cells tested from species such as humans and rabbits (7). Both viral DNA replication and late protein synthesis are significantly inhibited causing abortive infection during Δ*M062R* infection, while early gene expression remain intact (7). The virotherapeutic benefit we observed prompted us to investigate the immunological effect caused by Δ*M062R* (18), since this mutant virus is unable to establish a productive viral replication and oncolytic effect of Δ*M062R* is not through direct induction of cell death, e.g., apoptosis. We performed a RT^2^ profiler screening for antiviral responses to compare how responses generated by Δ*M062R* infection differed from the wildtype MYXV infection. We found Δ*M062R* infection in human primary monocytes stimulated the expression of many antiviral and interferon-stimulated genes (ISGs) (**Figure 2A**). To validate the above findings from the screening we collected periphery blood from 4 healthy individuals and purified CD14^+^ monocytes/macrophages for RT^2^-PCR confirmation. We found Δ*M062R* infection in these cells generally stimulated elevated IFN-I and ISGs, e.g., IFN β and CXCL-10 (**Figure 2B**), consistent with the screening results. We also confirmed the elevated CXCL-10 levels in the supernatant of Δ*M062R* infected cells (**Figure 2C**). As a control we included a MYXV deletion mutant in which another *C7L* superfamily gene, *M063R*, was ablated, *M063R*-null MYXV (19). *M063R*-null MYXV remains replication-competent in human cells and *M063R* does not possess a broad-spectrum host range function (17, 19). In our study, the control *M063R*-null MYXV infection did not induce IRF-dependent luciferase expression (not shown), and similar to the wildtype virus *M063R*-null MYXV did not cause upregulation of CXCL10 in human CD14^+^ macrophages (**Figure 2C**).

**Figure 2.**
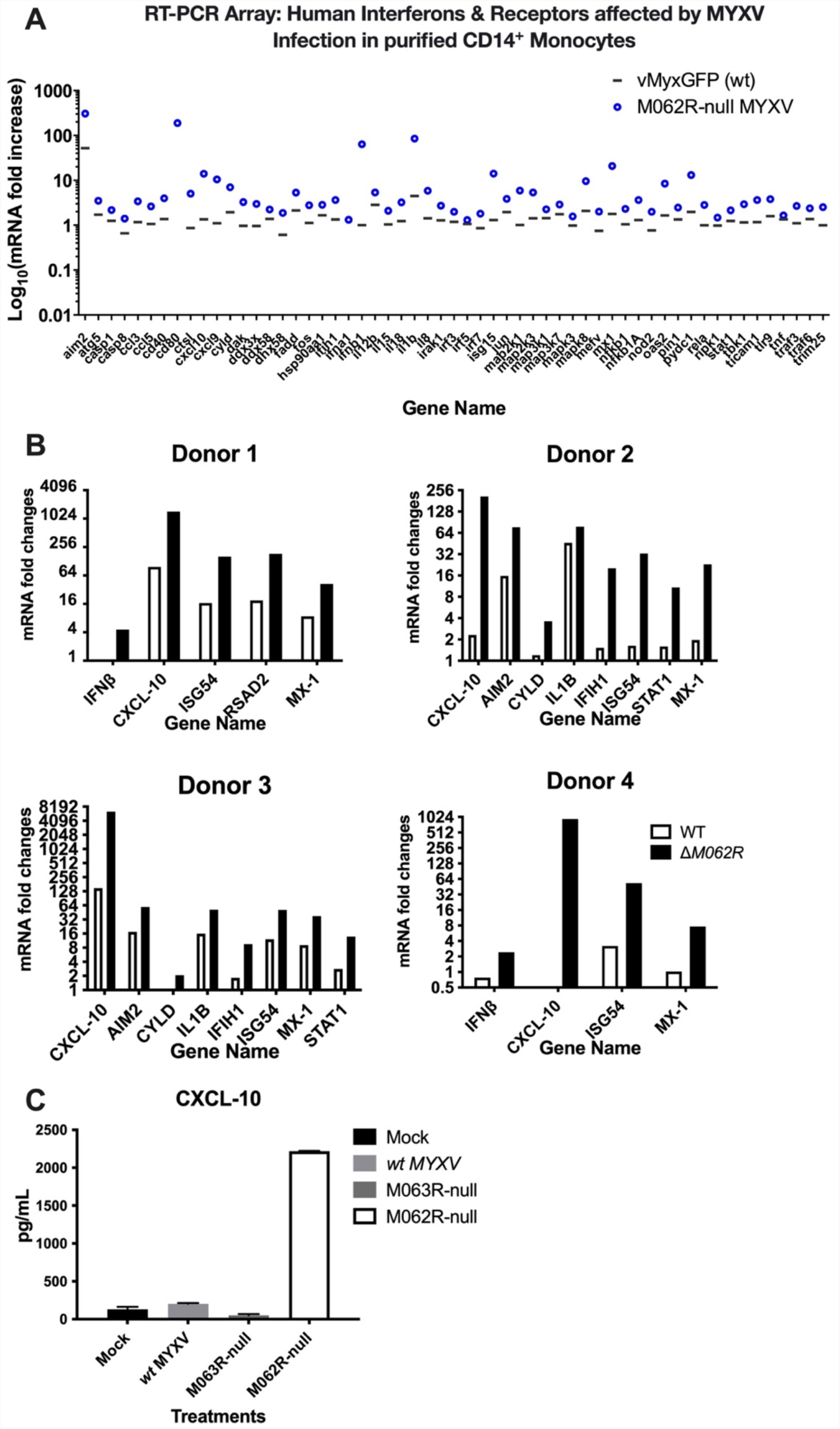
Infection by Δ*M062R* MYXV stimulates the expression of interferon-stimulated genes (ISGs). **A**. Infection by Δ*M062R* MYXV stimulates higher levels of IFN and ISG RNAs in primary human CD14^+^ monocytes than that from the wildtype MYXV infection. Infection by MYXV, wildtype and Δ*M062R*, was performed at an moi of 10 and cells were collected at 16 hours post-infection (p.i.) for RNA extraction, reverse transcription, and RT-PCR according to the manufacturer’s protocol. The ΔCt and ΔΔCt was calculated and RNA levels were normalized to an internal control gene, B2M. By normalizing to the result of the mock infected group for the same gene, fold induction of the corresponding gene is calculated using the formula of Foldchange=2^(-ΔΔCt). Shown is representative data from one of two healthy donors. **B**. Validation of RT^2^-PCR by ΔM*062R* leads to upregulation of IFN-I and pro-inflammatory molecules in CD14. Primary CD14^+^ human monocytes from 4 healthy human donors were mock infected or infected with either wildtype or Δ*062R* MYXV. In each case the expression of mRNAs encoding several ISGs including CXCL-10, ISG54 and MX-1 were elevated in the monocytes/macrophages infected with ΔM*062R* MYXV compared to monocytes/macrophages infected with wild type MYXV. Fold changes are measured by normalizing to that of mock infection. **C**. Infection by ΔM*062R* can no longer inhibit DNA-sensing pathway. As a control for viral manipulation the *M063R*-null MYXV was used to infect primary CD14^+^ human monocytes along with other controls (mock infection and wildtype MYXV) and ΔM*062R* infection. CXCL-10 production in the supernatant is shown and the Δ*M062R*-infected monocytes/macrophages secreted significantly higher levels of CXCL-10 than control groups.

### The Δ*M062R* stimulated IRF-dependent gene expression is sensed through cGAS

We have reported previously that the infection defect by Δ*M062R* was due to its inability to overcome host SAMD9 function (8). However, the immunological impact of Δ*M062R* remains unknown. A computational analysis of SAMD9 across all homologues found putative DNA binding domains (20), and DNA pulldown experiments using either the VACV 70mer dsDNA (21) or herring testes dsDNA (not shown) indicated that SAMD9 was co-immunoprecipitated with dsDNA (**Figure 3A**). We found previously that the MYXV M062 protein binds to amino acids (aa) 1-385 of human SAMD9 but not to the first 285 aa residues or c-terminal portion of SAMD9 (18), while the region of 285-385 aa in human SAMD9 overlaps with the putative DNA binding domain, the Alba-2 domain (20). We next examined if a human SAMD9 1-385 aa fragment could also be associated with dsDNA. We used the *SAMD9*-null HeLa cells we previously engineered (18) for the experiment and by transiently transfecting the cells to express SAMD9 1-385 aa we performed dsDNA pulldown experiment similar to what is shown in Figure 3A. We found that the SAMD9 1-385 aa truncated protein also associated with dsDNA. As a control, we transiently expressed a SAMD9 N-terminal fragment of 1-110 aa that contains the SAM domain in *SAMD9*-null cells, and found that this SAMD9 fragment was not associated with DNA. We then investigated whether the expression of MYXV M062 might interfere with SAMD9’s presence in the dsDNA pulldown content. As a control, we infected HeLa cells expressing intact endogenous SAMD9 with either wildtype MYXV or Δ*M062R*. Wildtype MYXV infection significantly reduced the amount of SAMD9 associated with dsDNA compared with that from Δ*M062R* infection (**Figure 3C**).

**Figure 3.**
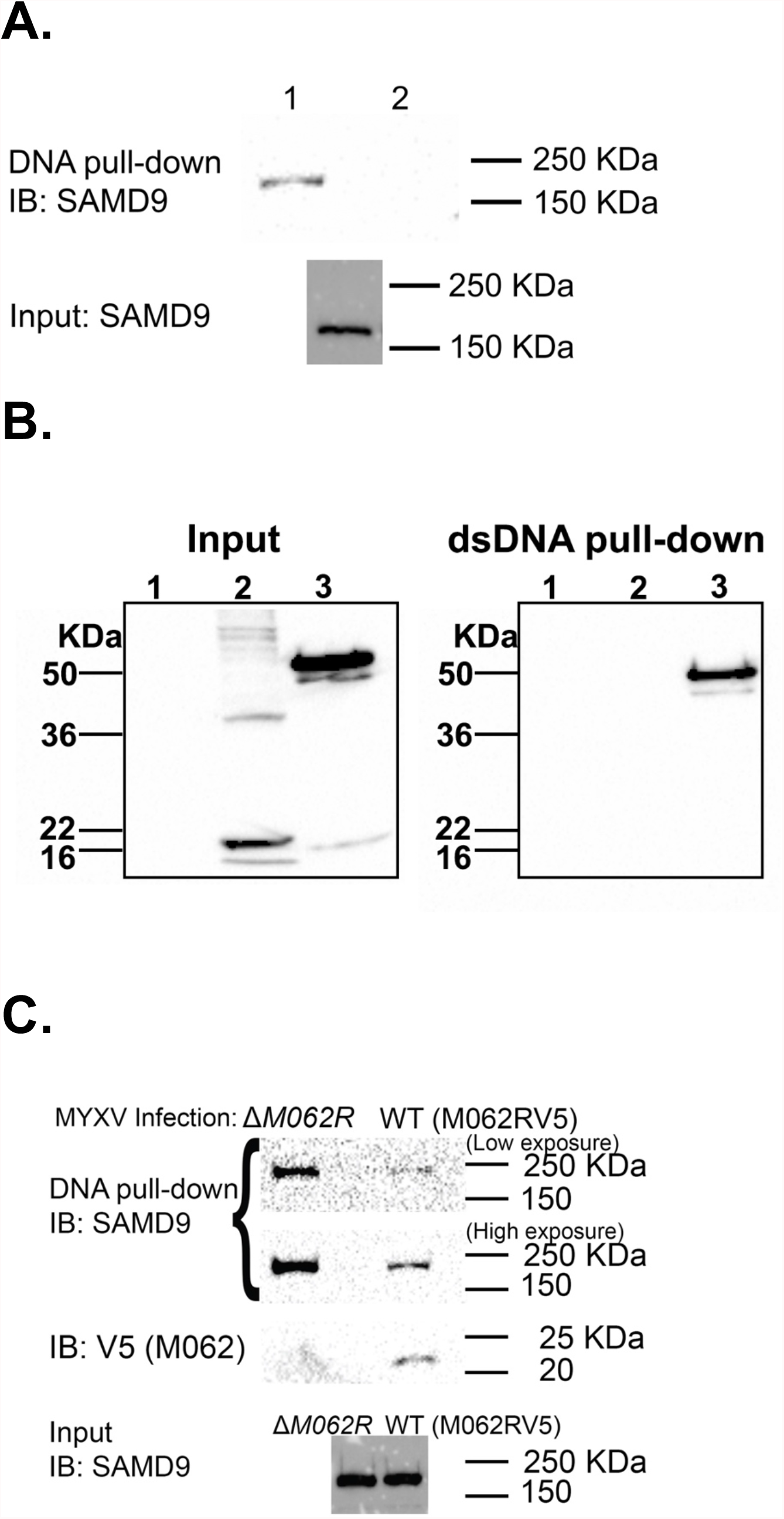
Viral M062 prevents human SAMD9 from being associated with dsDNA. **A**. DNA pull-down assay shows SAMD9 associated with dsDNA. HeLa cell lysate was incubated with either pre-conjugated Streptavidin UltraLink Resin with 5’-biotinylated VACV 70mer dsDNA or un-conjugated Streptavidin Resin alone. After extensive washing, resin-associated content was eluted for Western blot analysis by probing for SAMD9. Lane 1: 5’-biotinylated VACV 70mer dsDNA with resin; Lane 2: resin alone. **B**. SAMD9 1-385 aa domain but not the N-terminal 1-110 aa is associated with dsDNA. The same 5’-biotinylated VACV 70mer dsDNA pull-down experiment as in “A” is performed. Cell lysate from mock transfected, 1-110 aa construct (FLAG tagged) transfected, or 1-385 aa (FLAG tagged) construct transfected cells were used to incubate with dsDNA (VACV 70mer) conjugated resin and the co-precipitated content was separated on SDS-PAGE and probed for FLAG by Western Blot. **C**. The presence of M062 inhibits the association of SAMD9 with dsDNA. With the same dsDNA probe described above, we used HeLa cells expressing endogenous SAMD9 and infected them with either wildtype MYXV expressing V5 tagged M062 protein or ΔM*062R* MYXV for dsDNA pull-down assay. Proteins associated with DNA were separated on SDS-PAGE for Western Blot probing for SAMD9 and V5 tagged M062. The input lysates were examined for total SAMD9 expression.

Considering that SAMD9 may function through forming a complex with factors binding to DNA, we decided to test whether the ability of Δ*M062R* to induce IFN-I is due to the activation of DNA sensors. We utilized a luciferase expression in human monocytic THP-1 cells for the study. THP-1 cells can be differentiated into macrophages for testing DNA sensing and downstream outcome, and the firefly luciferase (F-Luc) expression is driven by the IRF recognition domain (Invivogen, San Diego, CA) (22). To test whether DNA sensing plays a role in Δ*M062R*-induced IFN-I induction and pro-inflammatory responses, we used cGAS-null THP-1 cells that were engineered from the F-Luc expressing parental cells mentioned above (22). We found that Δ*M062R* mutant virus stimulated robust luciferase expression comparable to that induced by interferon-stimulating DNA (ISD) (21) transfection (**Figure 4A**). In the absence of cGAS, the luciferase expression caused by both Δ*M062R* and ISD was eliminated (**Figure 4A**). However, transfection of 2’3’-cGAMP, the messenger molecule generated by cGAS upon DNA binding, successfully bypassed the lack of cGAS in cGAS-null THP-1 to restore F-Luc expression (**Figure 4B**). We thus conclude that the immunostimulatory effect of Δ*M062R* is due to the activation of cGAS-dependent DNA sensing pathway.

**Figure 4.**
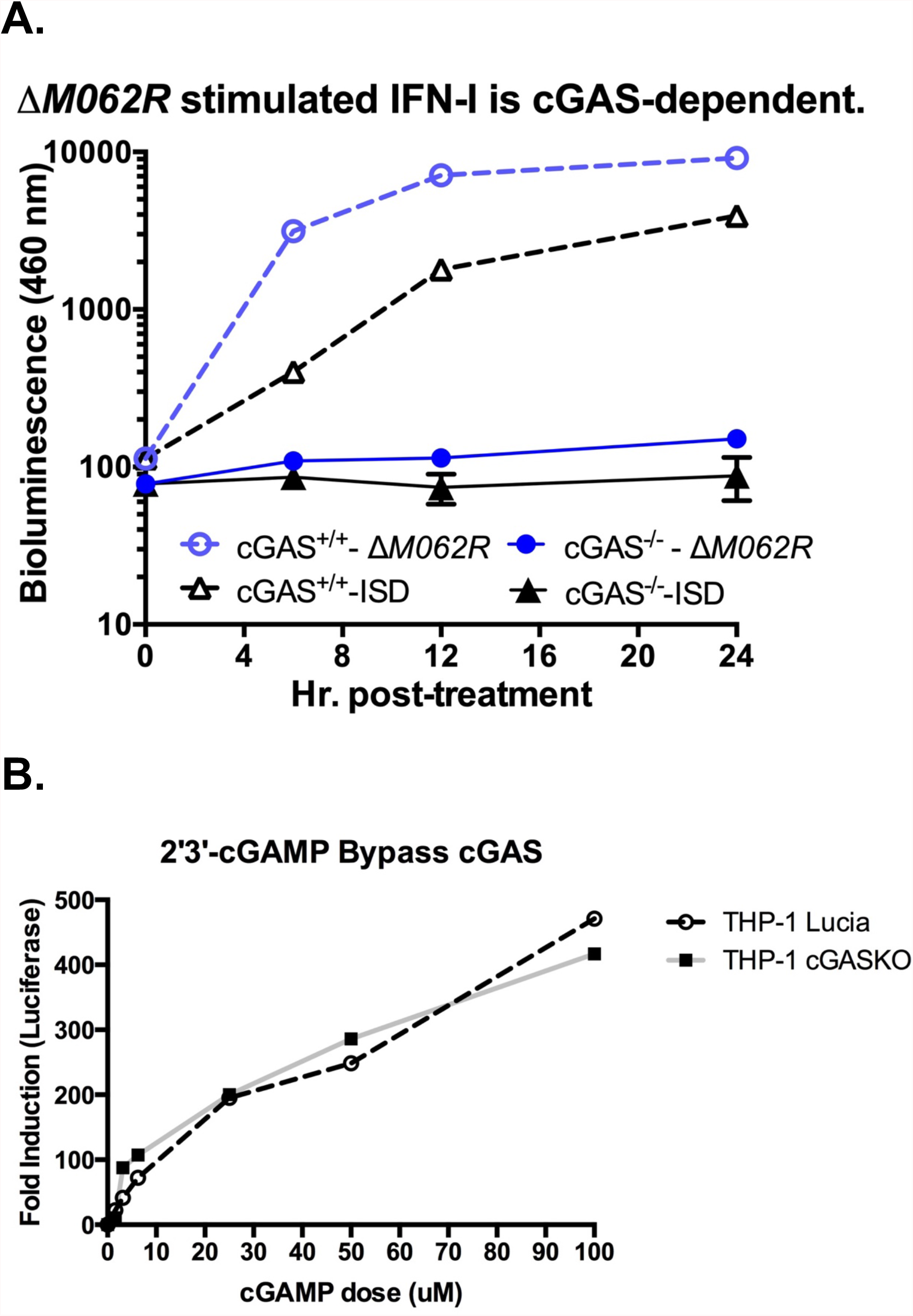
IRF-dependent gene expression stimulated by Δ*M062R* MYXV is regulated through cGAS. **A**. Infection by ΔM*062R* MYXV stimulates similar IRF-dependent gene expression to dsDNA-stimulated effect and is cGAS-dependent. THP-1-Lucia human macrophages with or without cGAS expression were treated by transfection with interferon-stimulated dsDNA (ISDs) or by infection with Δ*M062R* MYXV. Only the THP-1 cells with intact cGAS were able to respond with increased production of luciferase in either treatment. **B**. Addition of 2’3’-cGAMP bypasses the cGAS deficit and induces the production of IFN-I messenger RNA by RT-PCR. Transfection of 2’3’-cGAMP, the enzymatic product from activated cGAS, produces an identical response in both wild type and cGAS-null THP-1 macrophages, demonstrating that the remainder of the DNA-stimulated IFN-I pathway remains intact.

### Knocking down SAMD9 expression in monocytes/macrophages attenuated their proinflammatory responses

MYXV M062 inhibits SAMD9 function, leading to a productive viral infection. We next examined whether SAMD9 played a role in regulating the proinflammatory responses induced by Δ*M062R*. We generated stable SAMD9 knock-down THP1 cells using lentivirus expressing shRNAs targeting human SAMD9. As control, we engineered THP1 cells stably expressing scrambled shRNAs. We infected differentiated THP1 control or SAMD9 knockdown cells with Δ*M062R* for 18 hrs before examining pro-inflammatory cytokine production via RT-PCR. We found that reduced SAMD9 expression indeed attenuated Δ*M062R*-induced pro-inflammatory responses (**Figure 5A**). Transfection of ISD in these cells showed a similar attenuation in the IFNβ mRNA levels (**Figure 5B**). However, transfection of 2’3’-cGAMP led to upregulation of the IFNβ and ISG expression in SAMD9-knockdown THP1 cells (**Figure 5C**) that is similar to the response in the control THP1 cells. We thus concluded that Δ*M062R* infection stimulated a unique pro-inflammatory state that is cGAS-dependent and also regulated by SAMD9.

**Figure 5.**
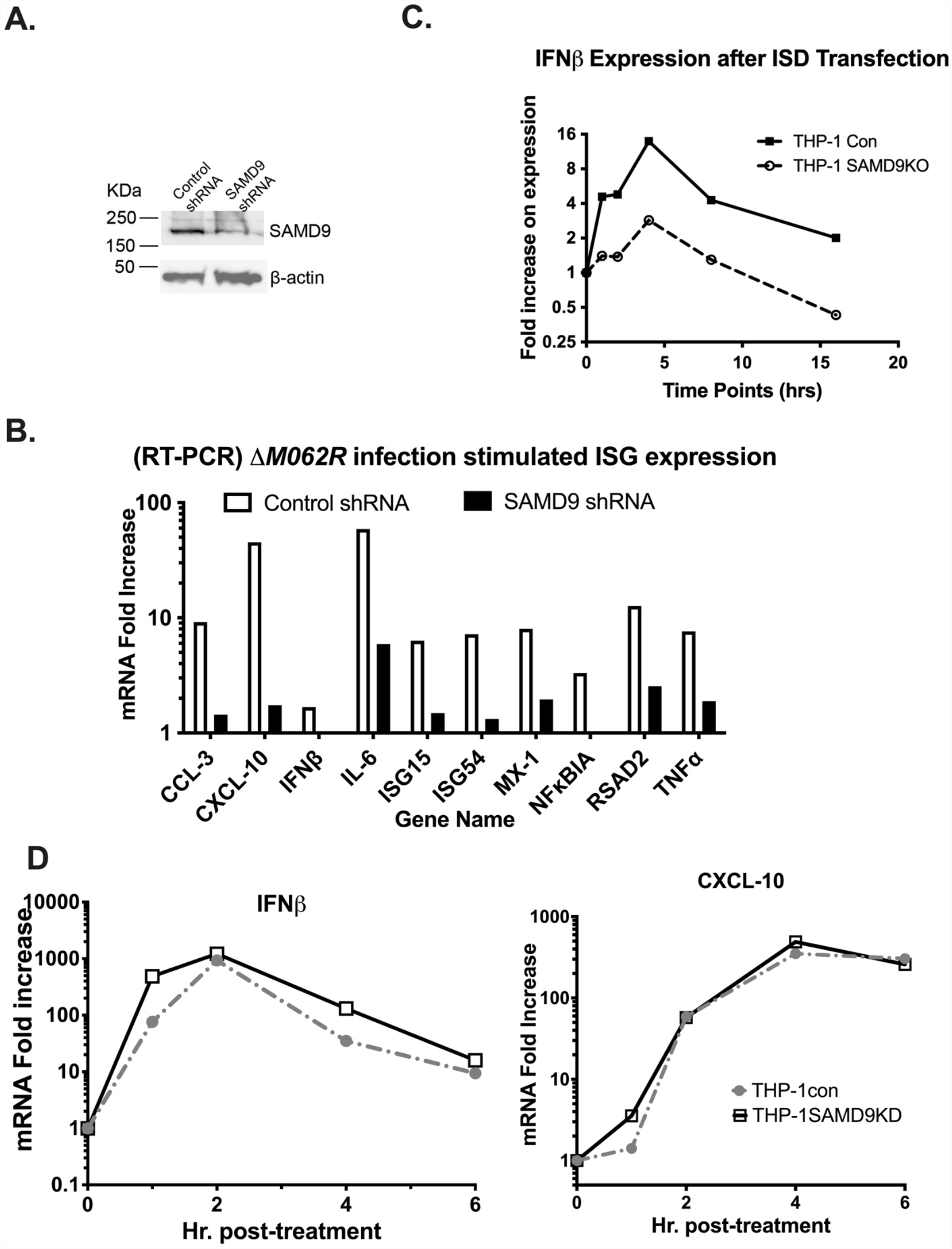
Knocking down SAMD9 expression attenuates the proinflammatory responses in THP-1 cells induced by Δ*M062R* infection. **A**. THP-1 cells in which SAMD9 was knocked down by shRNA. Western blotting showed reduced SAMD9 expression in THP-1 cells stably-transduced with lentivirus expressing SAMD9 shRNAs. **B**. Knocking down SAMD9 expression attenuated the proinflammatory response induced by Δ*M062R*. Compared to control shRNA expressing THP-1, the SAMD9-knockdown cells showed reduced proinflammatory responses after Δ*M062R* infection using RT-PCR. **C**. ISD-stimulated IFNβ mRNA is reduced in SAMD9 knockdown cells. After ISD transfection, IFNβ mRNA is measured by RT^2^-PCR and in SAMD9-knockdown cells the expression level of IFNβ is reduced. **D**. Transfection of 2’3’-cGAMP bypasses the block on dsDNA-induced proinflammatory responses in SAMD9-knockdown cells. Transfection of 2’3’-cGAMP in both control and SAMD9-knockdown cells showed similar expression of IFNβ and CXCL-10 mRNA.

### Next generation sequencing showed a unique gene expression profile during Δ*M062R* infection

We conducted a next generation sequencing study in the macrophage-like THP-1 cells to investigate the global transcriptomic change caused by Δ*M062R*. We hypothesized that since Δ*M062R* infection of macrophages stimulated cGAS-dependent IFN-I response, the infection will lead to similar results as dsDNA stimulated changes at the transcriptomic landscape. As control, we included the cells transfected with ISD dsDNA. Using the dual RNAseq bioinformatic analyses, we found that Δ*M062R* infection in monocytes/macrophages stimulated a very different gene expression profile from that of the ISD group (PCA plot **Figure 6A**, Venn diagram **Figure 6B**, and heatmap **Figure 6C**). Ingenuity Pathway Analysis (IPA) confirmed the activation of cGAS pathways by Δ*M062R* infection (IPA graphic summary **Figure 6E**), but additional gene expression profiling suggests a unique alteration of the transcriptomic landscape unlike that of the ISD transfection-induced antiviral response (**Figure 6F**). At the transcriptomic level, Δ*M062R* treatment also showed distinct changes compared to wildtype MYXV infection (**Figure 6D**). Although wildtype viral infection inhibited IRF3-dependent gene expression, it stimulated an unusual subgroup of chemokine/cytokine production in the macrophages (IPA graphic summary **supplemental Figure 1**).

**Figure 6.**
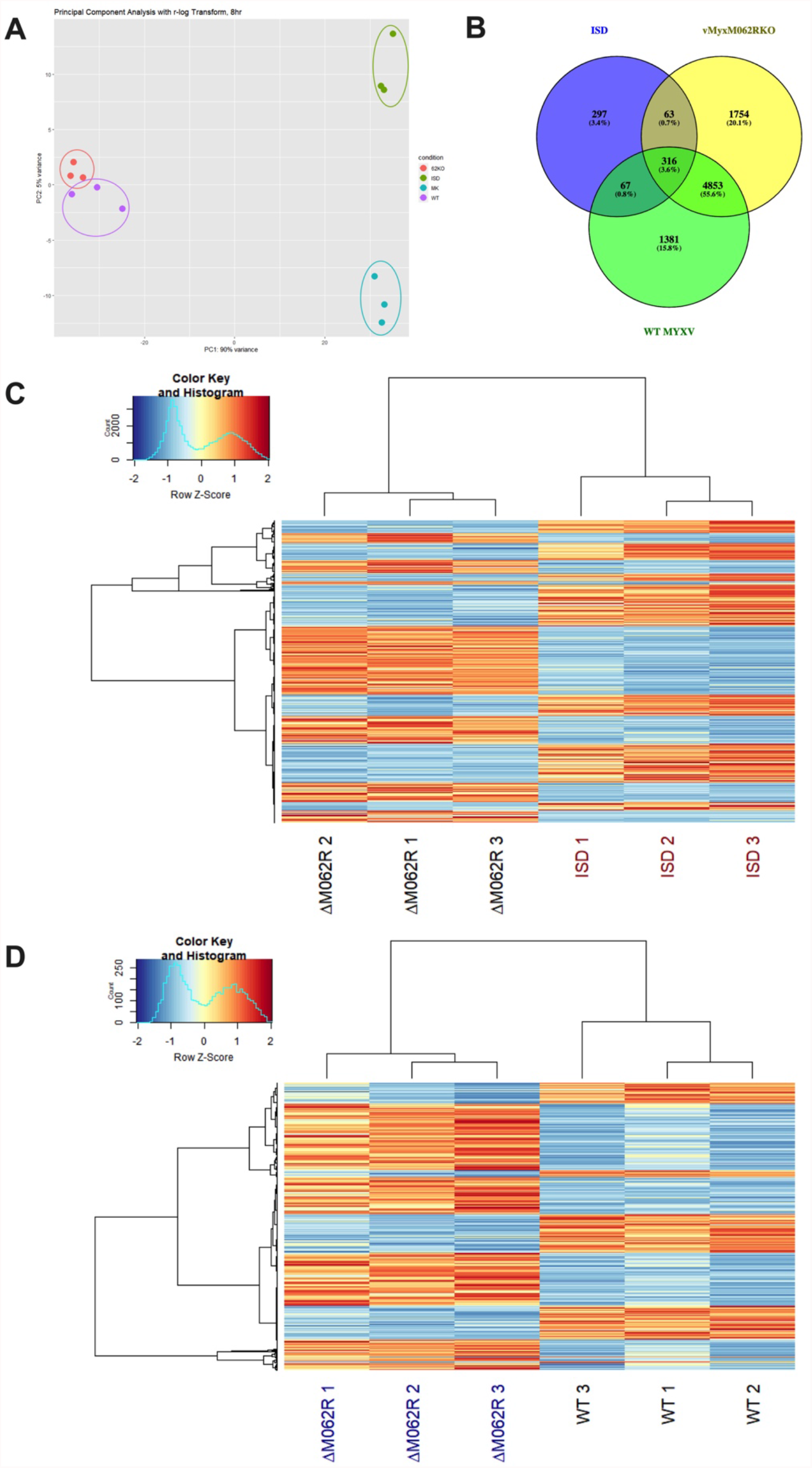
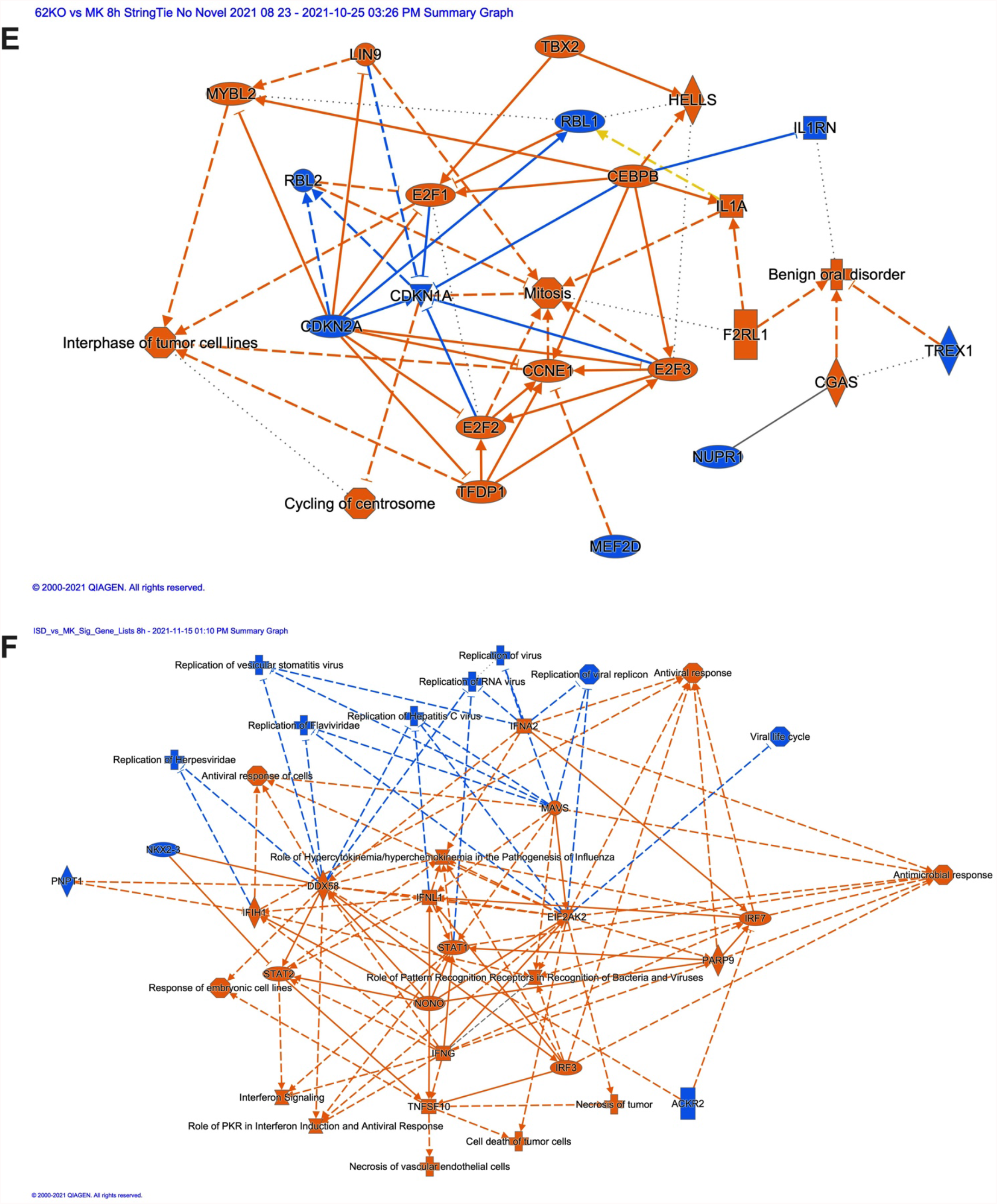
Dual RNAseq analyses reveal a unique host transcriptional profile stimulated by Δ*M062R* MYXV infection that is distinct from that caused by dsDNA. **A**. PCA plot of mock, ISD, ΔM*062R*, and wildtype MYXV treated samples. Good separation among all four samples are observed with close clustering within replicates. **B**. Venn diagram of host genes identified as being differentially regulated in ISD, ΔM*062R*, and wildtype MYXV groups compared to mock treated cells. Gene lists were generated by performing differential gene expression analysis using the R library DESeQ2, and only those genes whose adjusted-p-value was less than 0.1 were included in the analysis. **C**. Heatmap showing distinct host gene expression profile stimulated by Δ*M062R* from that by dsDNA. All significant differentially expressed genes between 2 groups under study were visualized using the R library heatmap.2. Gene counts were normalized using DESeQ2’s default normalization. Each row’s values were scaled using a z-score method before plotting and ward. D hierarchical clustering was performed using the Euclidean distance measure. **D**. Distinct host transcription profiles between ΔM*062R* and wildtype MYXV infection. All significant differentially expressed genes, between the groups under study, were visualized using the R library heatmap.2. Gene counts were normalized using DESeQ2’s default normalization. Each row’s values were scaled using a z-score method before plotting and ward. D hierarchical clustering was performed using the Euclidean distance measure. **E**. Ingenuity pathway analysis (IPA) revealed signaling pathway stimulated by Δ*M062R* infection. Host genes differentially expressed during Δ*M062R* infection at 8h post-infection were analyzed using IPA (Qiagen, version 01-20-04) for pathways affected. The graphic summary is shown. **F**. ISD transfection leads to activation of classic antiviral responses consistent to dsDNA-stimulated signaling events. Shown is the graphic summary generated by IPA using differentially expressed genes from ISD treatment group at 8-hour time point.

We next compared the viral gene expression pattern at 8 hours post-infection between Δ*M062R* mutant virus and wildtype MYXV. Surprisingly, in the Δ*M062R* infection we detected all the viral genes expressed in the wildtype MYXV infection (**Supplemental Figure 2**). We found 57 viral genes in Δ*M062R* infection were differentially expressed compared to wildtype MYXV with statistical significance (p value ranging from 1.03e-03 to 2.72e-88). Most of them (52 viral genes) (**Table 2**) showed slightly higher levels in Δ*M062R* infection than that in wildtype MYXV infection. *M136R* expression in the Δ*M062R* infection is 2.6-fold (logFC=1.4) higher than that expressed in wildtype MYXV infection (**Table 2**). Only 5 viral genes showed significant reduction at RNA levels during Δ*M062R* infection compared to that in wildtype MYXV infection. Among these viral genes, only *M062R* transcript was noticeably reduced (logFC=-3.95) due to the deletion of the central 80% of the *M062R* coding sequence (7) (**Supplemental Figure 3**) and the reduction in the levels of the remaining viral genes are in the range of 0.44-0.64 (logFC between −1.16 and −0.652) (**Table 2**).

## Discussion

Poxviruses are large dsDNA viruses with an exclusively cytoplasmic life cycle. These viruses can effectively inhibit IFN-I induction through many strategies. These strategies include blocking IFN-I signaling through decoy receptors (23), inhibiting key signaling effector molecules of the sensing pathways (9, 11), and/or reprograming host gene expression in the nucleus (24-26). One critical ability is for mammalian poxviruses, especially virulent poxviruses, to circumvent host immunosurveillance and the antiviral immune responses induced as a result of DNA sensing (9). Myxoma virus belongs to the genus of *Leporipoxvirus* and has a narrow host tropism, causing infectious disease only in rabbits. In European rabbits, virulent MYXV can cause lethal infection with 100% lethality and profound immunosuppression (27). Despite its limited host tropism for infectious disease, we found wildtype MYXV inhibits the DNA sensing pathway in human cells comparably to VACV, suggesting the presence of an antagonistic mechanism directed against host DNA sensing in a species-independent manner.

Host detection of cytoplasmic DNA by the cyclic GMP-AMP synthase (cGAS) leads to the production of the second messenger molecule 2’3’-cGAMP; binding of cGAMP to the adaptor, a stimulator of interferon genes (STING), triggers the activation of IFN-I production through a series of signaling events involving STING activation, recruitment and activation of TBK1, and phosphorylation of IRF3 (28). Many poxvirus proteins inhibit DNA sensing through different strategies, such as direct degradation of 2’, 3’-cGAMP by Poxvins (12, 13), inactivation of STING through mTOR by VACV F17 (15), inhibition of IRF3 by VACV C6 (29), and inhibition NF-κB activation by many VACV proteins including B14 (30) and F14 (31), etc. More importantly, inhibition of host DNA sensing by poxviruses during early infection seems to be a common strategy in spite of their species tropism. Poxvirus early and post-replicative gene expression can be distinguished through the use of a DNA replication inhibitor such as AraC. It is not surprising that poxviruses from different genera diverge on strategies to evade DNA sensing with distinct mechanisms but achieve the same outcome. In the MYXV genome, homologs of Poxvin are not found and a homolog of VACV F17 is predicted to be a late protein. MYXV may encode one or multiple additional early genes uniquely functioning as inhibitors of the DNA sensing pathway. In this study, we identified one MYXV viral protein which functions in such a capacity.

The MYXV *M062R* gene belongs to one of most conserved poxvirus host range factor families, the poxvirus *C7L* superfamily (17), and is a host tropism determinant of MYXV essential for viral replication (7). Although MYXV *M062R* can compensate for the function of the VACV *C7L* gene (32), it is predicted to have functional divergency from C7 and other *C7L* family members of orthopoxviruses (17). The MYXV M062 is unique from C7 in that [a] in VACV the *C7L* gene is non-essential and [b] only when *C7L* and another orthopoxvirus host range gene *K1L* are both deleted does the defect in host range tropism (comparable to Δ*M062R* MYXV) and replication deficiency become apparent (33). A known function of the MYXV M062 protein is to inhibit the function of the host protein SAMD9 (7, 8), but direct immunological impact of the MYXV M062 protein is not known. More importantly, the immunotherapeutic potential of *M062R*-null MYXV as an adjuvant for cancer therapy (6) further motivated our investigation of its immunostimulatory mechanism. Targeted deletion of *M062R* gene in the MYXV genome resulted in a mutant virus that maintained early gene expression at the protein levels but showed reduced DNA replication without late viral proteins being detected through western blot (7). Interestingly, in our RNAseq analyses we not only detected post-replicative viral RNA, especially late viral RNAs during Δ*M062R* infection, but also found Δ*M062R* viral RNA synthesis patterns in macrophages closely resembling that of the wildtype MYXV infection. We observed an occasional reduction in RNA levels among a few late genes during Δ*M062R* infection, and most of the Δ*M062R* viral transcripts were present at slightly higher levels than that in the wildtype MYXV infection at the same time point. The presence of significant levels of late RNA during Δ*M062R* infection is unexpected, as in order to synthesize late viral RNA comparable to the wild type virus level, many intermediate proteins must be produced *de novo*. This phenotype suggests a unique antiviral state stimulated by Δ*M062R* MYXV infection. In this state, viral protein synthesis, especially late protein production, is inhibited, which effect is coupled with inhibition of viral DNA replication. We speculate that the unique antiviral effect of inhibiting viral protein synthesis may be connected to the DNA sensing event. An alternative possibility is that the absence of M062 during Δ*M062R* infection may lead to translation deceleration of viral proteins until a complete stop, when a large quantity of late viral proteins are needed to complete the life cycle. In this alternative scenario, the DNA-trigger immune response reported may be caused by suboptimal levels of viral immunoregulatory proteins. This is, however, less probable because of the observed robust IRF-dependent gene expression triggered by Δ*M062R*, comparable to what is directly induced by dsDNA as shown in the luciferase assay.

Cytoplasmic sensing of DNA to trigger protective inflammation plays a key role in host antiviral defense (34, 35). The causes of cytoplasmic DNA may vary during the lifetime of a mammalian cell, such as improperly processed cellular DNA due to DNA repair or replication defect (36), and foreign DNA such as during viral infection (9, 37). Once triggered, DNA sensing induced IFN-I production and inflammation will lead to dramatic changes in the immunological milieu that may ultimately alter the immune responses profoundly. Thus, there must be additional control mechanisms to monitor and then regulate DNA-dependent IFN-I induction. There are known downstream host control mechanisms to fine tune IFN-I responses. Other than negative feedback cascade to restrict the duration and extent of IFN-I responses, e.g., SOCS, regulation of IFN-α receptor (IFNAR), and USP18 (38-40), intracellular signaling events and miRNAs can also perform such function (41). Upstream of IFN-I production, there are also regulatory measures to pattern recognition receptors (PRRs) and their adaptors, e.g., AKT to cGAS (42), TMEM120A to STING (43), and RNF138 to TBK1 (44). Our work provides evidence of a novel mechanism to fine tune the IFN-I response, which may operate through cellular translation regulation to PRR activation (**Figure 7**), and ultimately alters the global transcriptional landscape.

**Figure 7.**
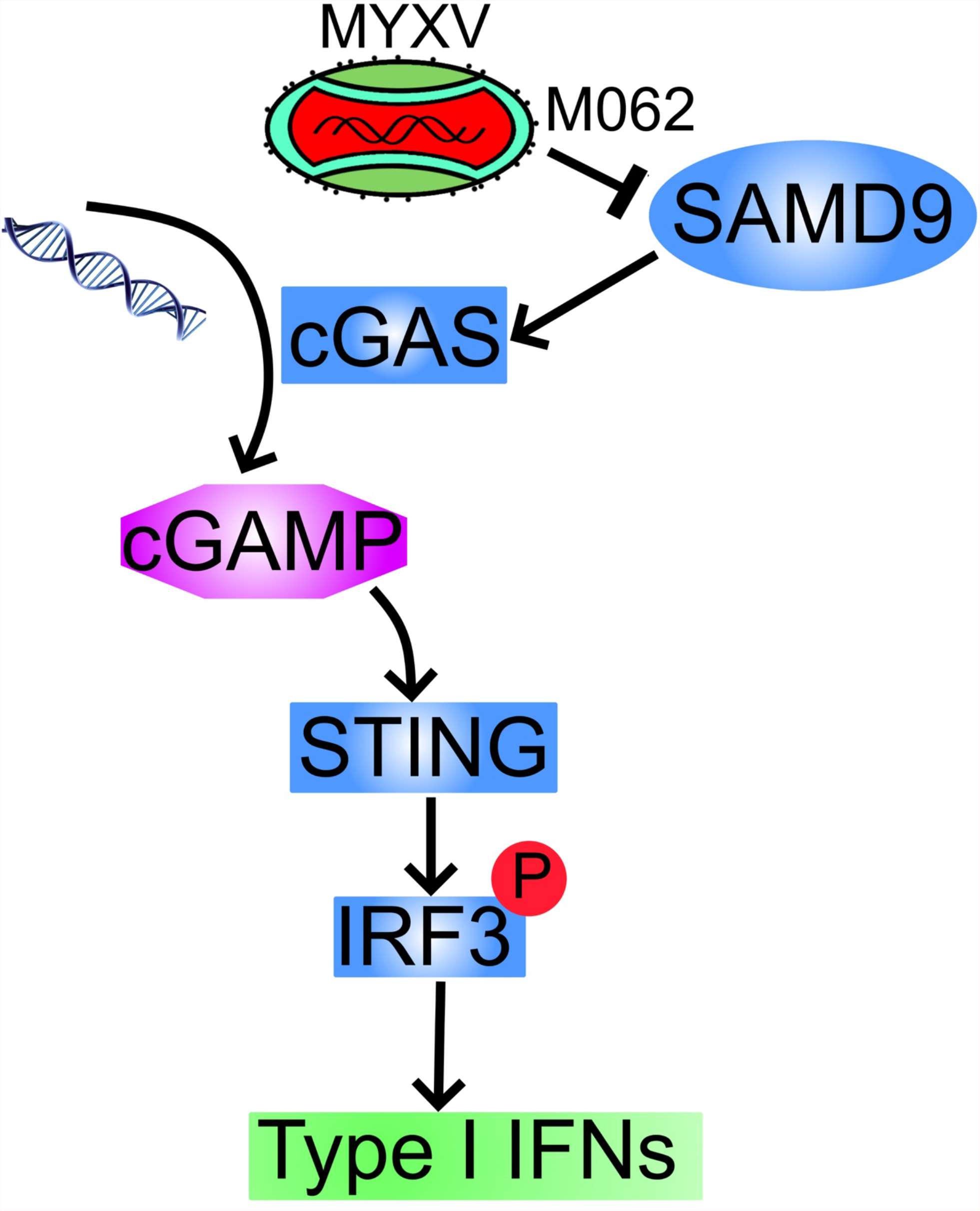
Summary of SAMD9 effect to DNA sensing pathway that is inhibited by viral.

SAMD9 is a large and exclusively cytoplasmic protein with complex domain structure, interestingly including putative DNA-binding domains (20). Our study indicated that SAMD9 itself is not necessarily a direct DNA sensor, since the triggering of IFN-I induction and subsequent proinflammatory responses is through cGAS-dependent DNA sensing events. During wildtype MYXV infection, cGAS may not have access to MYXV genome DNA in the cytoplasm in order to induce IFN-I. In the presence of M062 protein, SAMD9 is located exclusively outside of the cytoplasmic viral factories (8). However, during Δ*M062R* infection, SAMD9 can be detected abundantly surrounding the viral factories (8). Because of its putative DNA binding domain and its complex domain structure (20), SAMD9 may serve as an enabler to facilitate the access to DNA by sensors such as cGAS. SAMD9 participated in the regulation of host translation in the initiation and plays an important role in the elongation steps (unpublished data), and deleterious mutations in SAMD9 can inhibit global protein synthesis (45). More importantly, SAMD9 may serve as a signaling hub to fine tune the immunological consequences of the cell. SAMD9 may be targeted by other viruses (46) and has broad-spectrum antiviral effect (47-49). In our pursuit of understanding the mechanism of the SAMD9 antiviral effect, we revealed a key role of SAMD9 in regulating innate sensing for transcriptional remodeling in monocytes/macrophages. This work leads us to investigate its function in connecting innate immune sensing, translation control, and transcriptional refinement in host immune responses in the next step.

## Materials and Methods

### Cell culture and virus stock

Mammalian cells used in this study include BSC-40 (ATCC CRL-2761) (in Dulbecco minimal essential medium, DMEM, Lonza), healthy human periphery CD14^+^ monocytes (Lonza, Walkersville, MD), THP1 (ATCC TIB-202), and THP1-Lucia (kindly provided by F. Zhu) (50) (in RPMI1640, Lonza). The complete growth medium (e.g., DMEM Lonza/BioWhittaker Catalog no 12-604Q, or RMPI1640) was supplemented with 10% FBS (Atlanta Biologicals, Minneapolis, MN), 2 mM glutamine (Corning Cellgro, Millipore Sigma, St. Louis, MO), and 100 µ g per ml of Pen/Strep (Corning Cellgro, Millipore Sigma, St. Louis, MO); for RPMI1640 complete culture medium, in addition to FBS, glutamine, and Pen/Strep, 2-mercaptoethanol (MP biomedicals, Solon, OH) was supplemented to a final concentration of 0.05 µ M.

The viruses used were all derived from myxoma virus (MYXV), Lausanne strain (GenBank Accession AF170726.2). The MYXV *M062R* deletion mutant (vMyxM062RKOgfp) has been described previously (7). Myxoma virus stocks were prepared on BSC-40 cells and purified with sucrose step gradient through ultracentrifugation as previously described (51).

### RT^2^ Profiler PCR Array and Multi-plex Cytokine Array

Primary human CD14^+^ monocytes/macrophages (Lonza, Walkersville, MD) were mock treated, infected with wildtype MYXV and vMyxM062RKOgfp at a multiplicity of infection (moi) of 5 for 16 hours (hrs) before harvesting RNA for RT Realtime (RT^2^) PCR profiler and collecting supernatant for the Multi-plex Array study. For RT^2^ Profiler PCR Array Human Interferons and Receptors (Cat# PAHS064ZC-12, Qiagen, Germantown, MD), RNA extraction (RNeasy Pls Mini kit, Qiagen), cDNA synthesis (RT^2^ First Strand Kit, Qiagen), and Realtime PCR (RT^2^ SYBR Green qPCR Mastermixes, Qiagen) were performed following standard instructions from the manufacturer. The result is representative data from 1 individual and a total of 2 healthy individuals’ CD14^+^ monocytes which were tested (biological replicates). For Cytokine Multiplex of Human Inflammation Panel (Invitrogen™ Inflammation 20-Plex Human ProcartaPlex™ Panel) (Catalog # EXP20012185901, eBioscience/Invitrogen) procedures were performed following manufacturer protocol and the plate was processed on BioRad Bio-Plex 200 system with Bio-Plex-HTF attachment (Bio-Rad, Hercules, CA). The result is the representative from one individual’s CD14^+^ samples and a total of 2 healthy individuals’ CD14^+^ monocytes were tested. Shown is the average intensity from duplicate samples (technical replicates).

### Purification of human periphery CD14^+^ monocyte/macrophage

Healthy human whole blood was collected by venous puncture into collection tubes (BD catalog# 364606, Vacutainer ACD Solution), approximately 8 mL of whole blood per tube. An equal volume of sterile, room-temperature Dulbecco’s Phosphate Buffered Saline without calcium and magnesium (DPBS) (Corning catalog# 21-031-CV) was added to the flask. The diluted whole blood was layered over Lymphoprep solution (Accurate Chemical and Scientific Corp., Catalog# 1114545) and centrifuged at 2,500 rpm for 20 minutes. The collected PBMCs were 1:1 diluted with DPBS. The tube was then centrifuged at 1500 rpm for 5 minutes and the cell pellet was resuspended in 20 mL DPBS before repeating the wash process once more. The pellet was resuspended in 0.5 mL DPBS with 0.5 % BSA and labeled with 50 μL anti-CD14 microbeads (Miltenyi Biotec, catalog# 130-050-201). Cells were labeled for at least 20 minutes at 4 ° C and CD14^+^ cells were isolated by magnetic column (Miltenyi Biotec, catalog# 130-042-201).

### Luciferase assay

Supernatant from each sample was collected and immediately used in the luciferase assay. Luciferase presence in the supernatant was quantified by kit (QUANTI-luc, InvivoGen, catalog number rep-qlc1). Each sample was tested in triplicate in white 96-well plates with clear bottoms (LUMITRAC 200, Greiner Bio-One, Monroe, NC) with 5 times of volume to the supernatant sample. For each sample in triplicate the arithmetic average was reported. Fold induction is calculated as previously reported (9).

### Semi-quantitative RT Realtime PCR

THP-1 cells (10^6^ cells per 3.5 cm dish) were differentiated in PMA at 50 ng/mL for 48 hours (hrs) before being mock treated, transfected with ISD (21) at 2 µ g per dish using ViaFect (Promega, Madison, WI) at 3 µ L per 1 µ g of ISD based on manufacturer protocol, or infected at an moi of 5 for either WT (vMyxGFP) or *M062R*-null MYXV (7). At 1 hour (h) and 8 hrs post-transfection for ISD or 1 hr and 12 hrs post-infection, cells were harvested with Direct-zol RNA Mini Prep kit (catalog # R2052, Zymo, Irvine, CA) according to manufacturer standard protocol. RNA quality is examined by running on the RNA gel to check 28S and 18S integrity and spectrophotometer to estimate concentration. Equal amount of total RNA in a maximal volume of 6 µ L is used for cDNA synthesis using NEB PhotoScript® First Strand cDNA Synthesis Kit (Catalog # E6300L, NEB Inc, Ipswitch, MA) as instructed in the manufacturer standard protocol. Realtime PCR is conducted following manufacturer standard protocol (Luna Universal qPCR Master Mix, NEB Inc). Sybr green RT-PCR primers used in this study is listed in Table 1.

**Table 1.**
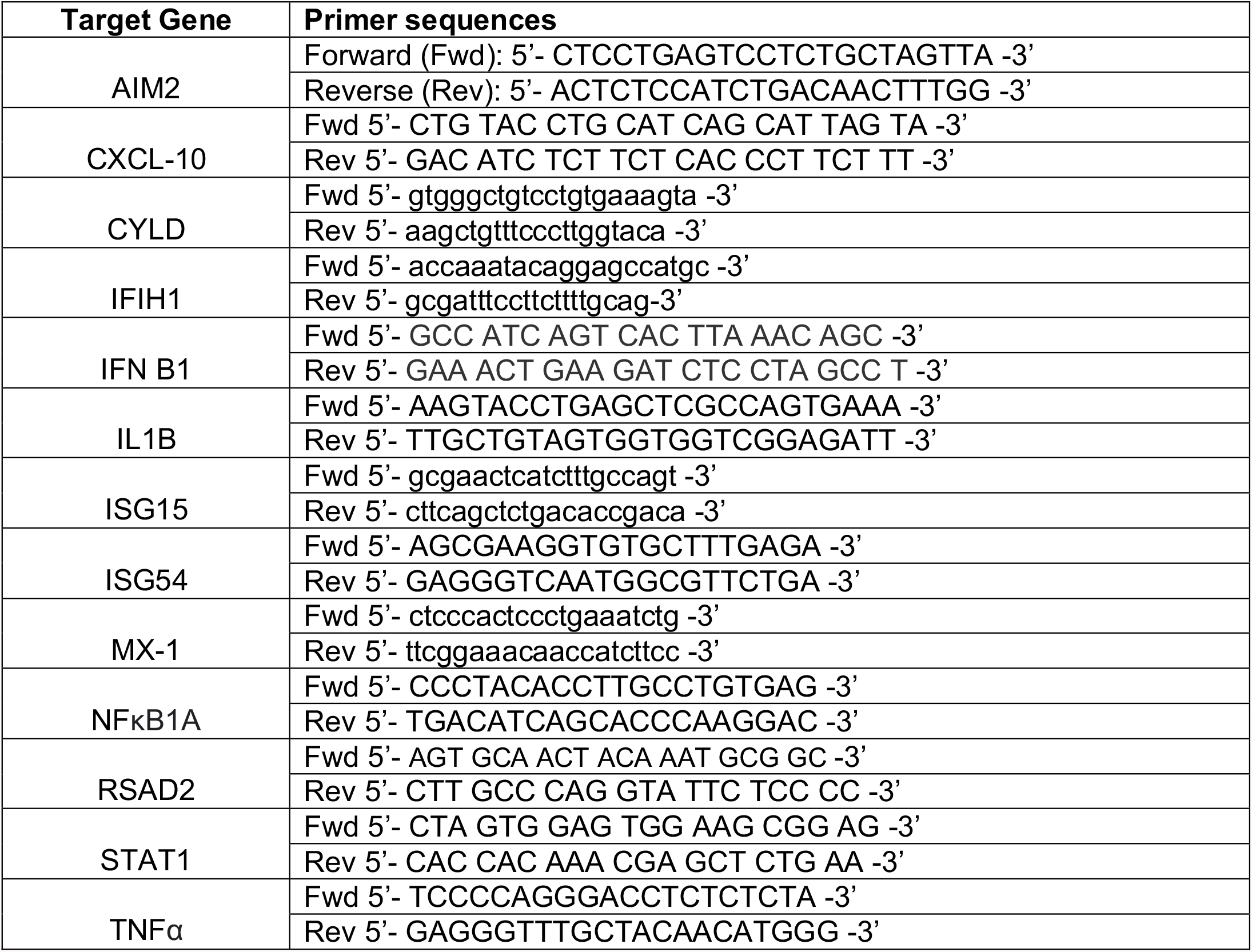
Real-time PCR Primer sequences.

**Table 2.** Differentially expressed viral genes in Δ*M062R* and wildtype MYXV in human macrophages.

| Uniprot ID | Gene ID | Expression kinetics | logFC | p-Value | Conservation within poxviruses |
| --- | --- | --- | --- | --- | --- |
| MYXV_gp005 | M003.2L | Unknown | 0.8 | 1.34e-03 | Sermi-conserved |
| MYXV_gp007 | M004.1L | Late | 0.801 | 3.22e-03 | Unique |
| MYXV_gp008 | M005L (M-T5) | Early | 0.835 | 9.9e-05 | Sermi-conserved |
| MYXV_gp009 | M006L | Early | 0.928 | 6.18e-6 | Sermi-conserved |
| MYXV_gp011 | M008L | Early | 0.6980 | 7.88e-05 | Sermi-conserved |
| MYXV_gp012 | M008.1L | Late | 0.901 | 8.94e-06 | Uniuqe |
| MYXV_gp015 | M011L | Early | 0.637 | 8.45e-04 | Sermi-conserved |
| MYXV_gp017 | M013L | Early | 0.489 | 4.18e-03 | Unique |
| MYXV_gp022 | M018L | Early | 0.544 | 4.39e-03 | Sermi-conserved |
| MYXV_gp023 | M019L | Late | 0.59 | 3.57e-03 | Conserved VACV F9L |
| MYXV_gp028 | M024L | Early | 0.883 | 4.32e-06 | Sermi-conserved |
| MYXV_gp031 | M027L | Late | 0.93 | 1.97e-06 | Conserved VACV E1L |
| MYXV_gp032 | M028L | Late | 0.617 | 4.87e-04 | Conserved VACV E2L |
| MYXV_gp035 | M031R | Early | 0.429 | 1.77e-03 | Sermi-conserved |
| MYXV_gp036 | M032R | Late | 0.604 | 1.78e-04 | Conserved |
| MYXV_gp038 | M034L | Early | 0.67 | 3.94e-04 | Conserved |
| MYXV_gp053 | M049R | Early | 0.39 | 5.72e-03 | Sermi-conserved |
| MYXV_gp055 | M051R | Unknown | 0.54 | 1.18e-03 | Sermi-conserved VACVG6R |
| MYXV_gp057 | M053R | Intermediate | 0.746 | 1.27e-04 | Conserved VACV G8R |
| MYXV_gp058 | M054R | Late | 0.824 | 2.44e-05 | Sermi-conserved VACV G9R |
| MYXV_gp059 | M055R | Late | 0.643 | 5.12e-04 | Conserved VACV L1R |
| MYXV_gp062 | M058R | Late | 0.507 | 1.77e-03 | Conserved VACV L4R |
| MYXV_gp066 | M062R | Early/Late | -3.95 | 2.72e-88 | Conserved VACV C7L |
| MYXV_gp067 | M063R | Early/Late | 0.56 | 5.56e-04 | Sermi-conserved VACV C7L |
| MYXV_gp072 | M068R | Early or Intermediate | 0.416 | 5.82e-03 | Conserved VACV J6R |
| MYXV_gp077 | M073R | Early | 0.6450 | 2.36e-04 | Sermi-conserved VACV H5R |
| MYXV_gp078 | M074R | Early/Late | 0.8280 | 1.5e-05 | Conserved VACV H6R |
| MYXV_gp079 | M075R | Early | 0.505 | 2.3e-03 | Sermi-conserved VACV H7R |
| MYXV_gp088 | M084R | Early | 0.774 | 3.37e-05 | Conserved VACV D9R |
| MYXV_gp106 | M102L | Late | -0.8610 | 4.51e-04 | Sermi-conserved VACV A13L |
| MYXV_gp107 | M103L | Late | -1.16 | 3.54e-05 | Conserved VACV A14L |
| MYXV_gp110 | M106L | Late | -0.6520 | 4.81e-04 | Conserved VACV A16L |
| MYXV_gp111 | M107L | Late | -0.8240 | 1.57e-03 | Sermi-conserved VACV A17L |
| MYXV_gp112 | M108R | Intermediate | 0.969 | 6.44e-06 | Conserved VACV A18R |
| MYXV_gp114 | M110L | Late | 0.802 | 9.72e-04 | Conserved VACV A21L |
| MYXV_gp119 | M115L | Late | 0.918 | 1.23e-06 | Sermi-conserved VACV A27L |
| MYXV_gp120 | M116L | Late | 1.02 | 2.7e-07 | Conserved VACV A28L |
| MYXV_gp121 | M117L | Early | 0.535 | 1.11e-03 | Conserved VACV A29L |
| MYXV_gp122 | M118L | Late | 0.678 | 2.5e-04 | Conserved VACV A30L |
| MYXV_gp123 | M119L | Early | 0.582 | 1.38e-03 | Unique |
| MYXV_gp126 | M122R | Late | 0.888 | 1.02e-04 | Sermi-conserved VACV A34R |
| MYXV_gp133 | M129R | Early | 0.615 | 1.37e-03 | Sermi-conserved VACV E7R |
| MYXV_gp134 | M130R | Early | 0.515 | 9.06e-04 | Unique |
| MYXV_gp140 | M136R | Late? | 1.4 | 5.54e-11 | Unique VACV A52R |
| MYXV_gp144 | M140R | Early | 0.599 | 1.06e-03 | Sermi-conserved VACV A55R |
| MYXV_gp145 | M141R | Early | 0.734 | 2.93e-04 | Unique |
| MYXV_gp148 | M144R | Early | 0.463 | 4.99e-03 | Sermi-conserved VACV N1L |
| MYXV_gp153 | M150R | Early | 0.565 | 1.03e-03 | Unique VACV C9L |
| MYXV_gp154 | M151R | Early | 0.792 | 9.40e-05 | Sermi-conserved VACV SPI-2 |
| MYXV_gp155 | M152R | Early | 0.843 | 5.60e-05 | Unique |
| MYXV_gp158 | M156R | Early | 0.638 | 2.51e-04 | Sermi-conserved VACV K3L |
| MYXV_gp159 | M008.1L | Late | 0.893 | 9.81e-06 | Unique |
| MYXV_gp160 | M008L | Early | 0.694 | 8.45e-05 | Sermi-conserved |
| MYXV_gp162 | M006R | Early | 0.931 | 5.51e-06 | Sermi-conserved |
| MYXV_gp163 | M005R (M-T5) | Early/Late | 0.833 | 1.05e-04 | Sermi-conserved |
| MYXV_gp164 | M004.1L | Late | 0.791 | 3.37e-03 | Unique |
| MYXV_gp166 | M003.2L | Unknown | 0.787 | 1.45e-03 | Sermi-conserved |

### Generation of SAMD9 knockdown and control THP1 cells

Lentiviral particles for stable SAMD9 shRNA expression (Cat# sc-89746-V, Santa Cruz Biotechnology, Dallas, TX) and Lentiviral particles for stable expression of scramble shRNAs (Santa Cruz Biotechnology, Dallas, TX) were used for generating the cell lines. We followed the manufacturer’s standard protocol similar to that reported before (7), and stable expression was selected under puromycin at 5 µ g/mL. SAMD9 knock-down was confirmed by Western Blot using rabbit anti-SAMD9 antibody (Cat# HPA021319, Millipore-Sigma, St. Louis, MO) and goat-anti-rabbit HRP secondary antibody (Cat# 111-035-144, Jackson ImmunoResearch Laboratory, INC, West Grove, PA) according to manufacturer’s instructions.

### Next Generation RNA sequencing

THP-1 cells (10^6^ cells per 3.5 cm dish) were differentiated in phorbol myristic acid (PMA) (Millipore-Sigma) at 50 ng/mL for 48 hours before mock treated, transfection with ISD (21) at 2 µ g per dish using ViaFect (Promega, Madison, WI) at 3 µ L per 1 µ g of ISD based on manufacturer recommendations, or infected at an moi of 5 for either WT (vMyxGFP) or *M062R*-null MYXV (7). At 8-hour post-transfection of ISD and at 1- and 8-hour post-viral infection cells were harvested, pelleted by centrifugation and then stored in −80° C until RNA extraction. RNA was extracted using the Quick DNA/RNA Mini-Prep Plus kit (catalog # D7003; Zymo, Irvine, CA, USA) with on-column DNase digestion. Purified RNA was assessed for mass concentration using the Qubit RNA BR Assay kit (catalog # Q10211; Invitrogen, Waltham, MA, USA) and for integrity using the Standard Sensitivity RNA Analysis kit on a Fragment Analyzer capillary electrophoresis system (catalog # DNF-471-0500; Agilent, Santa Clara, CA, USA). A total of 250ng total RNA was used for each sample as input to the TruSeq Stranded Total RNA library prep kit with unique-dual indexing (catalog #s 20020598 & 2002371; Illumina, San Diego, CA, USA). Libraries were assessed for mass using the Qubit 1X dsDNA HS Assay kit (catalog # Q33231; Invitrogen, Waltham, MA, USA), for fragment size using the High Sensitivity NGS Fragment Analysis kit on a Fragment Analyzer capillary electrophoresis system (catalog # DNF-474-0500; Agilent, Santa Clara, CA, USA), and functional validation using the Universal Library Quantification kit (catalog # 07960140001; KAPA, Wilmington, MA, USA).

Validated libraries were adjusted to 3nM before pooling, denaturing, and clustering. Paired-end (2×75) sequencing was performed to an average of 40 million reads per sample on a HiSeq 3000 (Illumina, San Diego, CA, USA).

### Dual RNAseq data processing and Ingenuity Pathway Analysis (IPA)

Raw Illumina binary base call (BCL) files were demultiplexed, adapter trimmed, and transformed to paired-end FASTQ files using bcl2fastq v2.18.0.12 (52). FastQC v0.11.4 (53) was then used to assess the quality of the FASTQ files. A “hybrid” *H. sapien* (Ensembl GRCh37 build) and Myxoma virus (Lausanne strain, NCBI, NC_001132.2) reference genome was constructed.

FASTQ files for each sample were aligned to the hybrid genome using STAR v2.7.6a (54) and run in its two-pass mode. STAR alignment metrics and QualiMap v2.2.1 (55) were used to assess alignment quality.

StringTie v2.1.4 (56) was then used to perform transcriptome reconstruction using each sample’s BAM file. Options were set to only allow the reconstruction and quantification of annotated genes. The gene reference general feature format (GTF/GFF) was produced. Additionally, options were provided to output “Ballgown-ready” files.

For gene-level exploratory data analysis (EDA) and differential expression analysis, StringTie output was imported into DESeQ2 v1.32.0 (57), where p-value and adjusted p-value thresholds were set to 0.05 and 0.1, respectively.

Differential gene expression analysis (e.g., *M062R*-null MYXV infection vs. mock and wildtype MYXV infection vs. mock) was uploaded for IPA to host pathway core analyses, and graphic summary was exported for data presentation.

### Statistical analyses

Graphpad Prism 9.1 was used for statistical analyses. Multiple-group comparison with single variable was performed using One-way ANOVA followed by secondary comparisons (e.g., Tuckey multiple comparisons test). Statistical significance is defined by a p<0.05.

## Acknowledgements

The study was supported by NIH K22AI099184 and R01AI139106 to J.L., a fund from the River Valley Ovarian Cancer Coalition, and a start-up by UAMS Department of Microbiology and Immunology to J.L. M.C. was supported by NIH P50 CA136393-11, Mayo Clinic SPORE in ovarian cancer. This work is also supported in part by the Center for Microbial Pathogenesis and Host Inflammatory Responses grant P20GM103625 through the NIH National Institute of General Medical Sciences (NIGMS) Centers of Biomedical Research Excellence (COBRE) and by the Winthrop P. Rockefeller Cancer Institute at UAMS. We would like to thank Jianmei Chen, Shana Chancellor, Pratikshya Paudel, and especially Sarah Blair for their technical support.

## Legends

**Supplemental Figure 1.**
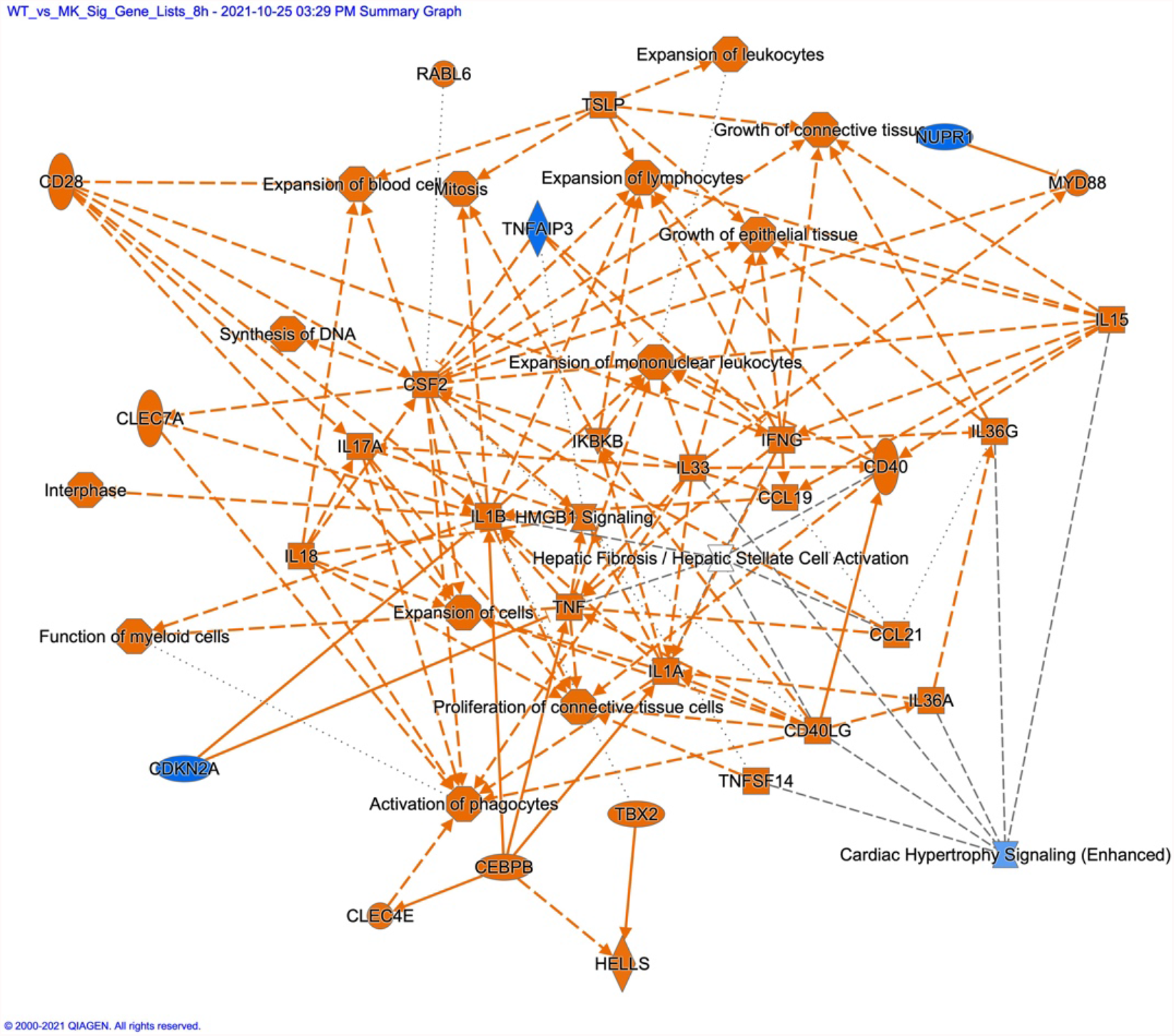
IPA analyses revealed unique transcriptional changes in host gene expression stimulated by wildtype MYXV. Host genes differentially expressed in wildtype MYXV infection were analyzed using IPA and a graphic summary of the results is shown.

**Supplemental Figure 2.**
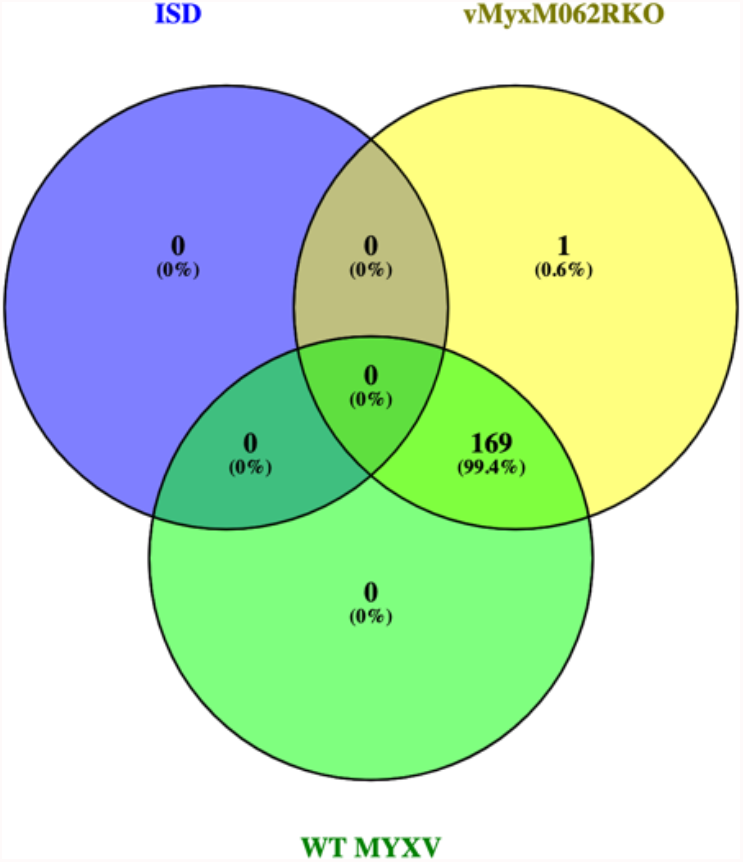
Venn diagram of MYXV specific genes detected in dual RNAseq analyses. Almost all viral genes expressed by wildtype MYXV at 8 h were detected in ΔM*062R* infection.

**Supplemental Figure 3.**
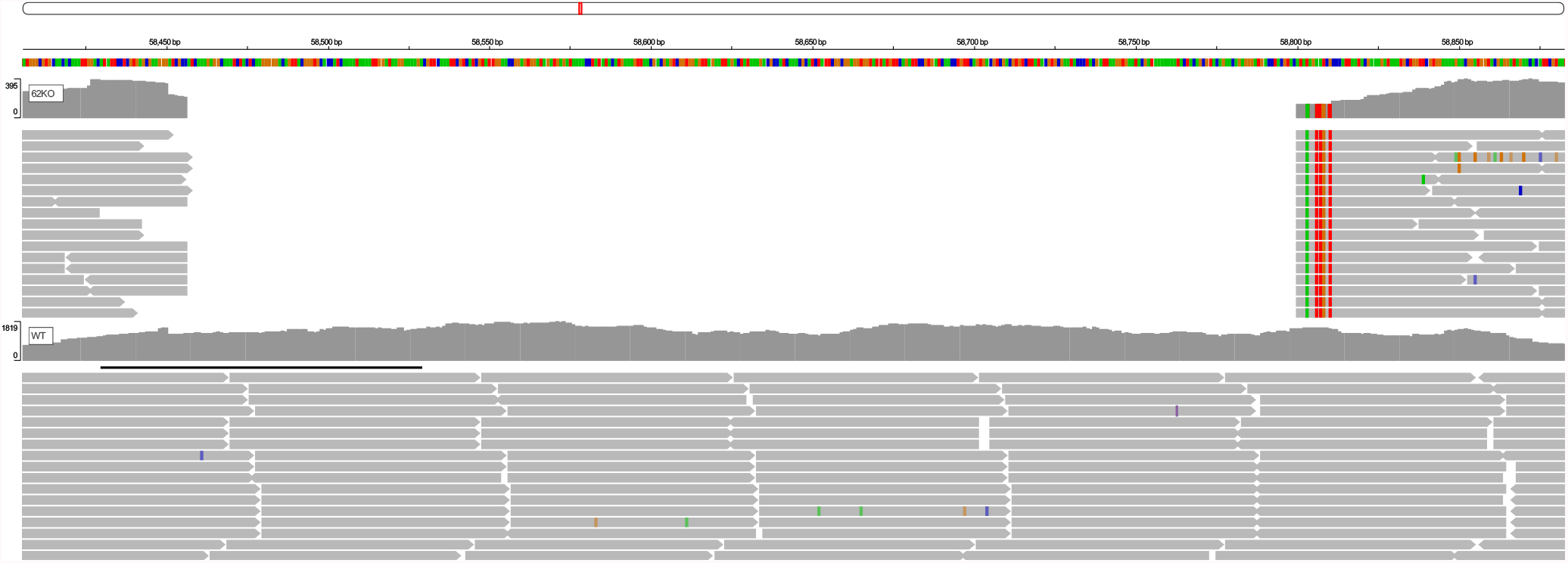
Raw data alignment of MYXV specific reads from Δ*M062R* and wildtype MYXV infection showed that only the *M062R* gene was absent in samples infected with the Δ*M062R* recombinant. Alignment files (i.e., BAM) from the first replicate from each of the 8 h Δ*M062R* and wildtype MYXV infection, was uploaded to igv.org/app/ for visualization. The dual genome reference was also uploaded and only the base pairs corresponding to the *M062R* gene were displayed.

